# Phage genome architecture and GC content: Structural genes and where to find them

**DOI:** 10.1101/2024.06.05.597531

**Authors:** Ritam Das, Janina Rahlff

## Abstract

Bacteriophages, the ubiquitous bacterial viruses, are the most abundant biological entities on Earth. However, information regarding the impact of environmental factors on the genetic makeup of phages is scarce. Based on the analysis of genomes of 45 phage isolates from different ecosystems we demonstrate a significant (*p* < 0.0001) positive correlation of the number of structural genes and their placement in above-average guanine-cytosine (GC) content of the genome. This feature appears to be more evident in isolates from deep-sea and sea-ice environments compared to desert and Baltic Sea slick, indicating a potential ecosystem-based factor (although insignificant due to low sample size). Additionally, the percentage of structural genes in above-average GC regions was found to be significantly (*p* = 0.0038) negatively correlated with the genome size of the phages and significantly (*p* = 0.0266) positively correlated with the overall GC for phage genomes > 50 kb, but not for genomes < 50 kb (*p =* 0.556). We therefore propose a relationship of structural genes and GC content, a property that may influence the adaptation and evolution of phages in various environmental niches.

## Introduction

Bacteriophages are proteinaceous biological entities composed of several structural elements which encapsulate, protect, and inject the genome into the bacterial host. This highly diverse group of virion proteins shelter the genetic material and abets in the recognition of the bacterial cell, playing a key role in viral replication (Cantu, *et al*. 2020).

GC content refers to the percentage of guanine (G) and cytosine (C) bases in a DNA or RNA molecule. The GC content is a crucial aspect of nucleic acid composition and can have significant implications for the structure, stability, and function of the genetic material.

In bacteria, high GC content has been positively associated with aerobiosis (Aslam, *et al*. 2019), nitrogen fixation (Mcewan, *et al*. 1998), and ultraviolet radiation from sunlight (Singer and Ames 1970). Certain bacterial groups are associated with a particularly low or high GC content often indicative to the ecosystem they inhabit. Marine SAR11 lineages and aquatic Actinobacteria feature low GC (Ghai, *et al*. 2013, Luo, *et al*. 2015), whereas Actinobacteria from soil or the stratosphere have a high GC content (Ellington, *et al*. 2021, Ghai, *et al*. 2011).

The GC content of viruses was shown to correlate with that of their hosts, even across kingdoms (Bahir, *et al*. 2009, Simon, *et al*. 2021). Such a positive correlation was also shown for bacteriophages and their hosts (Cardinale and Duffy 2011). Higher GC content was found in larger bacterial but smaller phage genomes (Almpanis, *et al*. 2018). Across 144 human DNA viruses, the GC content was lower in coding regions of small viral genomes compared to large genomes (Sewatanon, *et al*. 2007). High GC in the herpes simplex virus dsDNA was suggested to be a protection of viral genes against mobile genetic elements (Brown 2007). As for bacteria, potential ecosystem associations with the GC content of viruses have been observed. Viruses from marine metagenomes were found to have significantly lower GC content than viruses from the atmosphere (Rahlff, *et al*. 2023a). The genome of an induced prophage from the Arctic bacterial strain *Leeuwenhoekiella aequorea* Arc30, which has been isolated from the sea-surface microlayer, showed a clear placement of structural genes into regions of above-average GC within the genome. In contrast, genes related to DNA replication/recombination/repair and packaging genes were positioned in below-average GC regions (Rahlff, *et al*. 2024).

Here, we propose a relationship between the GC content and the position of structural genes in bacteriophage genomes and due to previous findings (see above) assume an association with overall viral GC content, genome size, and the ecosystem of the phage.

## Materials and methods

Genomes of phage isolates (n = 45, Table S1) were downloaded from NCBI’s GenBank and annotated using DRAM-v version 1.4.6 (Shaffer, *et al*. 2020). Using a sliding window of 10000 bp and step size of 100 bp within Proksee (Grant, *et al*. 2023), the GC content in relation to the location of the structural genes was determined. The overall GC content of the phage genomes was determined using emboss v. 5.0.0 (http://emboss.open-bio.org/). Data were checked for normal distribution using Shapiro-Wilk test, Kolmogorov-Smirnov test, and Anderson-Darling test in GraphPad Prism v.10. Depending on the outcome, either Pearson or Spearman rank correlation was applied. Ecosystem effects were investigated using the Kruskal-Wallis test with Dunn’s multiple comparisons. A list of the genes counted as structural genes is provided in Table S2.

## Results and discussion

The distribution of structural genes into above-average GC regions is representatively shown for two polar phage genomes (Fig. 1 a & b) and for two cases where the distribution is deviating from this architecture in a phage genome from a Baltic Sea surface slick and one from desert soil (Fig. 1 c & d). Across all phage genomes, the total number of phage structural genes in the genome was significantly and positively correlated to the number of structural genes in above-average GC regions (Pearson *r* = 0.6756, *n* = 45, *p* < 0.0001, Fig. 2A). The percentage of structural genes in above-average GC regions was significantly negatively correlated with genome size (Spearman *r* = -0.423, *n* = 45, *p* = 0.0038, Fig. 2C). It was significantly positively correlated with overall GC for phage genomes > 50 kb size (Pearson *r* = 0.611, *n* = 13, *p* = 0.0266, Fig. 2D) but not significantly correlated for phage genomes < 50 kb size (Pearson *r* = 0.108, n = 32, *p =* 0.556). When separating phages by the ecosystem (sea-ice, wastewater, marine, deep-sea, soil, haloalkaline lake, Baltic Sea slick, and desert) they have been isolated from, we observe insignificant trends (Kruskal-Wallis test) with desert and slick having the lowest, and deep-sea phages having on average the highest percentage of structural genes in above-average GC regions (Fig. 2B). We conclude, the feature is related to overall GC content, viral genome size and potentially to the ecosystem of the phage (with a possible relationship to temperature, hydrostatic pressure), the latter however needs further investigation using bigger sample size and a bigger variety of phages from different hosts. We speculate an above-average GC content of the structural genes could contribute to i) an increase in stability, bendability, and transcription of these genes (Vinogradov 2003) ii) the protection of the genetic material from thymine-dimer mediated UV damage (Nagpal, *et al*. 2021) iii) and/or in a broader sense assist the phage in resisting and replicating in extreme environments. For instance, in bacteria, DNA dynamics such as supercoiling might be more important during cold shocks than during heat shocks (Golovlev 2003).

**Figure 1:**
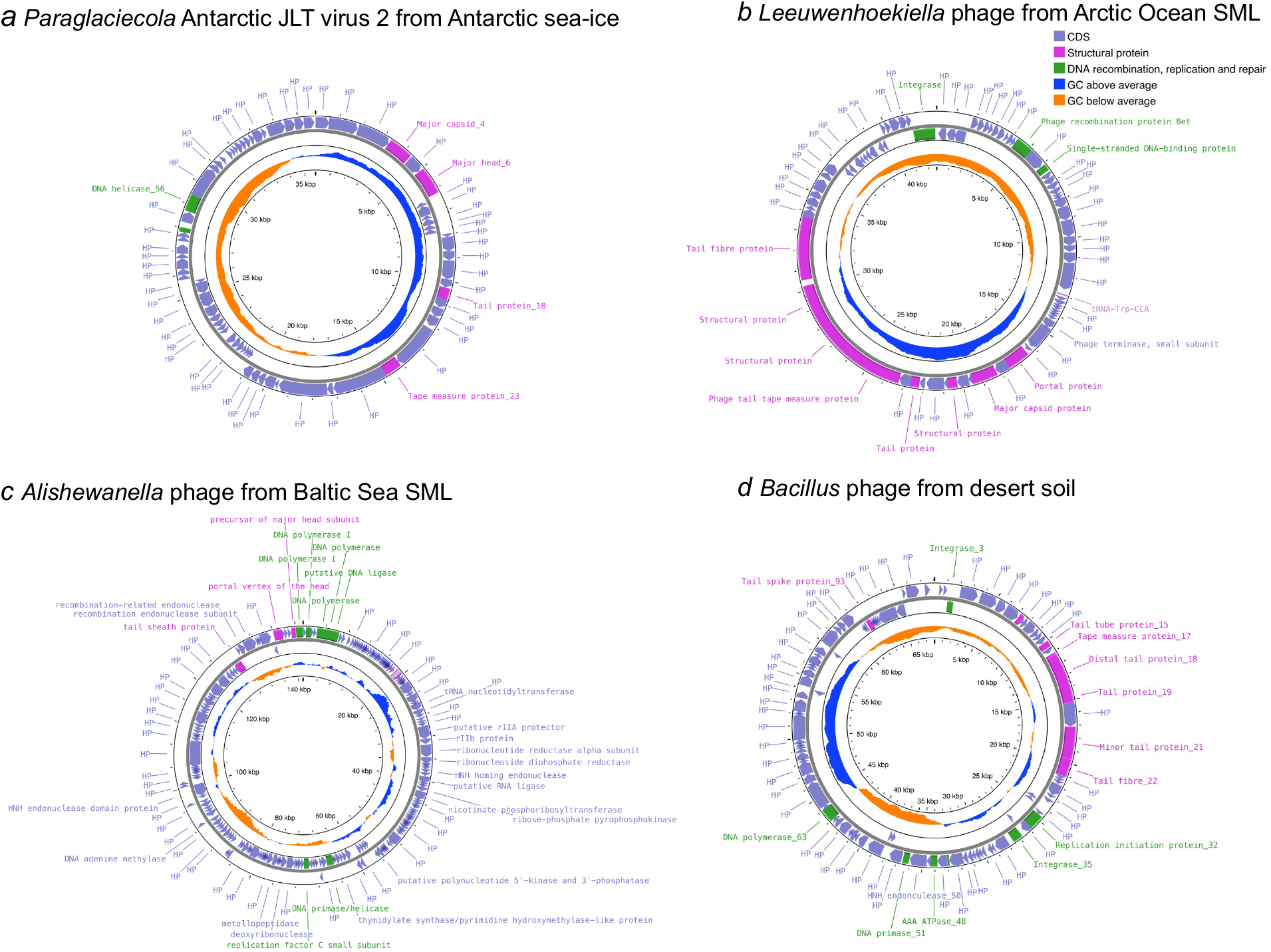
The architecture, GC patterns and functional annotations of four phage genomes: a) from *Paraglaciecola* Antarctic sea-ice virus (GenBank accession MW805362.1 (Demina, *et al*. 2022)), b) *Leeuwenhoekiella* phage from Arctic Ocean SML (Rahlff, *et al*. 2024), c) *Alishewanella* phage from Baltic Sea SML (GenBank accession OQ508956.1 (Rahlff, *et al*. 2023b), and d) a *Bacillus* phage from desert soil (GenBank accession ON210834.1 (Krukonis, *et al*. 2022)).

**Figure 2:**
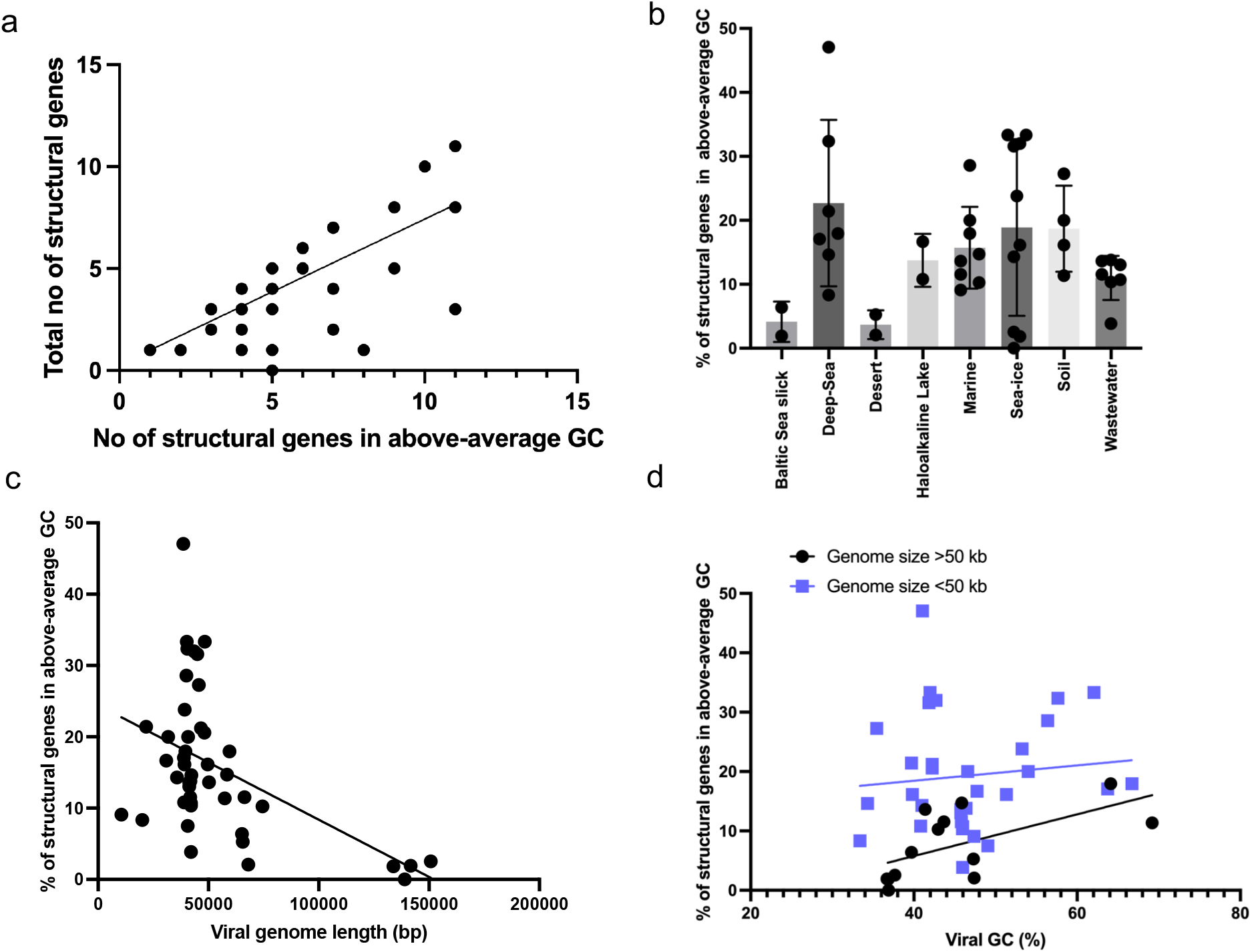
Correlations and bar plot for structural genes in above-average GC regions. The total number of all structural genes was positively correlated to the number of structural genes in above-average GC regions, b) the percentage of structural genes in phage genomes shown as separated by ecosystem, c) the negative correlation of the percentage of structural genes with genome length, and d) the relationship of percentage of structural genes in phage genomes with overall viral GC content for two different categories of genome size.

### Note

A poster with the same title has been presented at the International Virus Bioinformatics Meeting (ViBioM) 2024 in Leuven, Belgium. All annotations file are provided in the supplement.

## Supporting information

Annotation data

## Acknowledgements

RD received funding by German Research Foundation (DFG) grant “NFDI4Microbiota” number NFDI 28/1. JR received funding by the DFG Walter-Benjamin Return Grant RA3432/1-3, project number 534276621, and the Swedish Research Council, Starting Grant ID 2023-03310_VR. Data handling was enabled by resources provided by the National Academic Infrastructure for Supercomputing in Sweden (NAISS) and the Swedish National Infrastructure for Computing at UPPMAX partially funded by the Swedish Research Council through grant agreement no. 2022-06725.

## Supplementary Information

**Table S1:** List of phage genomes analyzed in this study.

| Phage | Accession ID | Length | GC% | Host | Ecosystem | Location | Reference |
| --- | --- | --- | --- | --- | --- | --- | --- |
| **P2559S** | JQ867099.1 | 44988 | 41.85 | *Croceibacter* | Sea-ice | Antarctica | https://doi.org/10.1128/jvi.01396-12 |
| **1/32** | KJ018210.1 | 42252 | 34.31 | *Flavobacterium* | Deep-Sea | Mariana Trench | https://doi.org/10.1111/1/1462-2920.12611 |
| **1/4** | KJ018209.1 | 133824 | 36.86 | *Shewanella* | Sea ice | Baltic Sea | https://doi.org/10.1111/1/1462-2920.12611 |
| **P12026** | JQ867100.1 | 31766 | 46.6 | *Marinomonas* | Sea | Yellow Sea | [10.1128/JVI.01328-12](https://doi.org/10.1128/JVI.01328-12) |
| **Antarctic GD virus 1** | MW805361.1 | 150766 | 37.67 | *Paraglaciecola* | Sea-ice | Antarctica | https://doi.org/10.1093/femsec/fiy028 |
| **Antarctic JLT virus 2** | MW805362.1 | 35731 | 40.98 | *Paraglaciecola* | Sea-ice | Antarctica | https://doi.org/10.1093/femsec/fiy028 |
| **Antarctic BD virus 1** | MW805363.1 | 48354 | 62.07 | *Octadecabacter* | Sea-ice | Antarctica | https://doi.org/10.1093/femsec/fiy028 |
| **Antarctic DB virus 2** | MW805364.1 | 39241 | 53.25 | *Octadecabacter* | Sea-ice | Antarctica | https://doi.org/10.1093/femsec/fiy028 |
| **268TH007** | ON210834.1 | 68062 | 47.38 | *Bacillus* | Tumamoc Hill Desert Laboratory | Tucson, Arizona | [10.1128/mra.00455-22](https://doi.org/10.1128/mra.00455-22) |
| **268TH002** | ON210835.1 | 65534 | 47.29 | *Bacillus* | Tumamoc Hill Desert Laboratory | Tucson, Arizona | [10.1128/mra.00455-22](https://doi.org/10.1128/mra.00455-22) |
| **vB_SmaS_Carrot** | OL539439.1 | 41293 | 45.74 | *Serratia* | Wastewater treatment plant (sewage) | Western United States | [10.1128/mra.01212-21](https://doi.org/10.1128/mra.01212-21) |
| **vB_SmaS_LittleDog** | OL539456.1 | 41738 | 45.81 | *Serratia* | Wastewater treatment plant (sewage) | Western United States | [10.1128/mra.01212-21](https://doi.org/10.1128/mra.01212-21) |
| **vB_SmaS_Niamh** | OL539455.1 | 42053 | 45.96 | *Serratia* | Wastewater treatment plant (sewage) | Western United States | [10.1128/mra.01212-21](https://doi.org/10.1128/mra.01212-21) |
| **vB_SmaS_Opt-148** | MW021766.1 | 41293 | 45.74 | *Serratia* | Wastewater treatment plant (sewage) | Western United States | [10.1128/mra.01212-21](https://doi.org/10.1128/mra.01212-21) |
| **vB_SmaS_Serratianator** | MW021755.1 | 42052 | 45.97 | *Serratia* | Wastewater treatment plant (sewage) | Western United States | [10.1128/mra.01212-21](https://doi.org/10.1128/mra.01212-21) |
| **vB_SmaS_Stoker** | OL539464.1 | 41797 | 46.38 | *Serratia* | Wastewater treatment plant (sewage) | Western United States | [10.1128/mra.01212-21](https://doi.org/10.1128/mra.01212-21) |
| **vB_SmaS_Ulliraptor** | OL539442.1 | 42052 | 45.97 | *Serratia* | Wastewater treatment plant (sewage) | Western United States | [10.1128/mra.01212-21](https://doi.org/10.1128/mra.01212-21) |
| **1/40** | KJ018211.1 | 139004 | 36.89 | *Shewanella* | Sea ice | Baltic Sea | https://doi.org/10.1111/1462-2920.12611 |
| **1/41** | KJ018212.1 | 43510 | 42.7 | *Shewanella* | Sea ice | Baltic Sea | https://doi.org/10.1111/1462-2920.12611 |
| **1/44** | KJ018213.1 | 49640 | 39.82 | *Shewanella* | Sea ice | Baltic Sea | https://doi.org/10.1111/1462-2920.12611 |
| **3/49** | KJ018214.1 | 40161 | 41.97 | *Shewanella* | Sea ice | Baltic Sea | https://doi.org/10.1111/1462-2920.12611 |
| **vB_PtuP_Slicky01** | OQ508958.1 | 65166 | 39.71 | *Pseudoalteromonas* | Slick | Baltic Sea | https://doi.org/10.1038/s43705-023-00307-8 |
| **vB_AspM_Slickus01** | OQ508957.1 | 141644 | 36.71 | *Alishewanella* | Slick | Baltic Sea | https://doi.org/10.1038/s43705-023-00307-8 |
| **vB_BhaS-171** | KU160496.1 | 38975 | 40.82 | *Bacillus* | Haloalkaline lake | Elmenteita, Kenya | [10.1371/journal.pone.0215734](https://doi.org/10.1371/journal.pone.0215734) |
| **vB_EauS-123** | KU160495.1 | 30924 | 47.73 | *Exiguobacterium* | Haloalkaline lake | Elmenteita, Kenya | [10.1371/journal.pone.0215734](https://doi.org/10.1371/journal.pone.0215734) |
| **vB_HmeY_H4907** | OQ832096.1 | 40452 | 57.64 | *Halomonas* | Deep-sea | Mariana Trench | https://doi.org/10.1128/spectrum.01912-23 |
| **vB_PaeM-G11** | OQ622254.1 | 40800 | 54.01 | *Pseudomonas* | Soil | Antarctic | [10.3390/ijms24087662](https://doi.org/10.3390/ijms24087662) |
| **vB_PeaS-P1** | KT381865.1 | 38868 | 63.75 | *Pelagibaca* | Deep-sea | No information | https://doi.org/10.1186/s12864-017-3886-0 |
| **vB_ThpS-P1** | KT381864.1 | 39591 | 66.73 | *Thiobacimonas* | Deep-sea | No information | https://doi.org/10.1186/s12864-017-3886-0 |
| **BVE2** | MG584725.1 | 20021 | 33.39 | *Bacillus* | Deep sea | Southwestern Indian Ocean | https://doi.org/10.1007/s00705-020-04579-6 |
| **Gxv1** | NC_049974.1 | 21781 | 39.71 | *Bacillus* | Deep sea | western Pacific Ocean | [10.1007/s42995-020-00074-8](https://doi.org/10.1007/s42995-020-00074-8) |
| **vB_BteM-A9Y** | ON528935.2 | 38634 | 41.05 | *Bacillus* | Deep sea | South China Sea | [10.3390/v15091919](https://doi.org/10.3390/v15091919) |
| **Waukesha92** | KJ920400.1 | 45648 | 35.45 | *Bacillus* | Soil | compost soil in Glenn Allen, Virginia | [10.1128/genomeA.00339-14](https://doi.org/10.1128/genomeA.00339-14) |
| **KBS-S-2A** | HQ634187.1 | 40658 | 49.07 | *Pseudanabaena* | Freshwater | Singapore Serangoon Reservoir | https://doi.org/10.1128/jvi.00682-20 |
| **DchS19** | ON287378.1 | 39149 | 51.34 | *Dickeya* | Soil | Sungai Petani, Kedah, Malaysia | [10.1128/mra.00800-22](https://doi.org/10.1128/mra.00800-22) |
| **vB_RpoS-V7** | MH015249.1 | 59573 | 64.12 | *Ruegeria* | Marine | Baltimore, MD | https://doi.org/10.1128/mra.00959-18 |
| **F115** | OP292673.1 | 48162 | 42.25 | *Salmonella* | Wastewater | Ecuador | https://doi.org/10.1128/mra.01048-22 |
| **F61** | OP292674.1 | 46635 | 42.25 | *Salmonella* | Wastewater | Ecuador | https://doi.org/10.1128/mra.01048-22 |
| **Abt2graduatex2** | MF975638.1 | 57385 | 69.19 | *Streptomyces* | Soil | Baltimore, MD | [10.1128/genomeA.01480-17](https://doi.org/10.1128/genomeA.01480-17) |
| **phiCB2047-B** | NC_020862.2 | 74485 | 42.98 | *Sulfitobacter* | Marine | Raunefjorden, Norway | [10.1128/genomeA.00945-13](https://doi.org/10.1128/genomeA.00945-13) |
| **phiGT1** | MT584811.1 | 40019 | 56.39 | *Sulfitobacter* | Marine | Yellow Sea, South Korea | [10.1128/MRA.00779-20](https://doi.org/10.1128/MRA.00779-20) |
| **CHOED** | KJ192399.2 | 66316 | 43.68 | *Vibrio* | Marine | Chilean mussels | [10.1128/genomeA.00091-14](https://doi.org/10.1128/genomeA.00091-14) |
| **fNo16** | MH730557.1 | 10594 | 47.36 | *Vibrio* | Marine | No information | [10.1128/MRA.00020-19](https://doi.org/10.1128/MRA.00020-19) |
| **Phriendly** | MN062185.1 | 50218 | 41.39 | *Vibrio* | Marine | South Padre Island shore | [10.1128/MRA.01096-19](https://doi.org/10.1128/MRA.01096-19) |
| **vB_VnaS-AQKL99** | MT795651.1 | 58464 | 45.87 | *Vibrio* | Marine | Bayboro Harbor | https://doi.org/10.1128/mra.00967-20 |

**Table S2:** List of genes counted as structural genes in this study.

| Sl. no. | Structural genes | Reference* |
| --- | --- | --- |
| 1 | Major head protein | https://doi.org/10.1371/journal.pcbi.1007845 |
| 2 | Major tail protein | https://doi.org/10.1371/journal.pcbi.1007845 |
| 3 | Tape measure protein | https://doi.org/10.1128/AEM.01236-12 |
| 4 | Tail protein | https://doi.org/10.1371/journal.pcbi.1007845 |
| 5 | Minor head protein | https://doi.org/10.1371/journal.pcbi.1007845 |
| 6 | Phage baseplate upper protein | https://doi.org/10.1371/journal.pcbi.1007845 |
| 7 | Tail spike protein | https://doi.org/10.3390/antibiotics11050555 |
| 8 | Tail completion protein | https://doi.org/10.1038/s42003-024-06221-6 |
| 9 | Phage tail tube protein | https://doi.org/10.3390/antibiotics11050555 |
| 10 | Tail fibre protein | https://doi.org/10.1371/journal.pcbi.1007845 |
| 11 | Tail terminator | https://doi.org/10.1016/j.jmb.2009.04.072 |
| 12 | Minor tail protein | https://doi.org/10.1371/journal.pcbi.1007845 |
| 13 | p22 coat protein | https://doi.org/10.3390/v6072708 |
| 14 | Head-tail adaptor | https://doi.org/10.1371/journal.pcbi.1007845 |
| 15 | Virion structural protein | https://doi.org/10.1093/gigascience/giac076 |
| 16 | Tail assembly protein | https://doi.org/10.3390/antibiotics11050555 |
| 17 | Tail completion or Neck1 protein | https://doi.org/10.1038/s42003-023-05292-1 |
| 18 | Tail sheath | https://doi.org/10.1371/journal.pcbi.1007845 |
| 19 | Portal protein | https://doi.org/10.1371/journal.pcbi.1007845 |
| 20 | Tail tip assembly protein | https://doi.org/10.1016/j.jmb.2013.03.032 |
| 21 | Bacteriophage HK97-gp10,<br>putative tail-component | PF04883, IPR010064 |

